# Use of anodal transcranial direct current stimulation for improving motor performance in healthy adults: A systematic review and meta-analysis

**DOI:** 10.64898/2026.05.01.722354

**Authors:** Atsushi Sasaki, Takuya Ideriha, Atsuya Matsuoka, Yujin Goto, Natsue Yoshimura, Nobuhiro Hagura

## Abstract

**Purpose:** Transcranial direct current stimulation (tDCS) can noninvasively modulate activity in targeted brain regions. It is well established that the excitability of motor-related regions can increase when the target region is located beneath the anode (anodal tDCS), suggesting its potential to increase motor performance. Although such attempts have been widely examined, the results remain inconclusive. The purpose of this study was to assess the conditions under which anodal tDCS may improve motor performance in healthy adults.

**Methods:** We conducted a systematic review of studies on the use of anodal tDCS for improving motor performance in healthy adults. A computerized search was performed using the Web of Science, Scopus, PubMed, JDreamIII, and Ichushi-Web to identify relevant studies published between January 1, 1990 and May 25, 2022.

**Results:** Twenty-five studies were included in the qualitative synthesis. For the meta-analysis, 25 trials (N=885) were extracted from 23 studies. There were significant effects of anodal tDCS on motor performance improvement, but with evidence of publication bias and substantial heterogeneity among the trials. Post-hoc analysis revealed that motor performance 24 hours after the application of anodal tDCS may benefit from stimulation. There was no marked effect related to stimulation intensity, duration, or whether stimulation was provided during motor performance.

**Conclusions:** Our study clarified the current state of anodal tDCS use for motor performance enhancement and indicates that there is currently no reliable evidence to support its effectiveness. Further studies, particularly randomized controlled trials, are necessary to establish the reliability of these effects for future applications.

## 1. INTRODUCTION

The use of noninvasive brain stimulation (NIBS) to enhance motor function is increasing [1]. Advancements in neuromodulation have expanded the potential applications of NIBS [2–4]. Among various NIBS techniques, transcranial magnetic stimulation (TMS) has been widely studied for its ability to modulate motor cortical excitability and improve motor function in individuals with neurological disorders [5,6]. Although TMS has demonstrated the ability to induce lasting changes in neuronal excitability [7], its practical limitations restrict its widespread application outside laboratory settings. Transcranial direct current stimulation (tDCS) is a more feasible and portable alternative for modulating brain activity [8]. With growing public and academic interest in tDCS [9], there is growing demand for clear evidence-based guidance regarding its use.

tDCS modulates neuronal excitability by delivering low-amplitude direct currents through scalp electrodes, producing polarity-specific effects: anodal stimulation is generally thought to increase cortical excitability, whereas cathodal stimulation decreases it [8,10]. These effects are believed to be mediated by alterations in the excitatory/inhibitory balance of local neural circuits, leading to changes in synaptic plasticity and, potentially, motor performance [10]. Based on these mechanisms, anodal tDCS over motor-related areas has been hypothesized to improve motor performance [8].

While many studies have employed NIBS techniques to transiently disrupt neural function to investigate causal brain–behavior relationships [11], the present study takes a different approach by focusing on the potential of tDCS to enhance motor performance in healthy individuals. This direction is particularly relevant, given the increasing application of tDCS outside laboratory settings, where expectations for performance enhancement are often high. However, despite widespread claims that tDCS improves various aspects of cognition, mood, and physical performance [12], the reproducibility and specificity of these effects remain controversial. Notably, some studies have raised substantial concerns regarding the robustness of tDCS-induced effects in healthy populations [13]. Although physiological measures suggest heightened corticospinal excitability following anodal tDCS [8], it remains unclear whether these neural changes translate into consistent performance benefits. Therefore, the uncritical use of tDCS in healthy users may lead to an overestimation of its practical utility or unintended interference with optimal motor function. Moreover, inappropriate use may raise safety concerns [14].

Several systematic reviews have reported encouraging evidence that anodal tDCS can improve selected aspects of motor performance [15,16]. These effects have been attributed to the tDCS-induced modulation of cortical excitability and its interaction with motor practice. However, despite these positive trends, findings across studies remain mixed. For instance, a review by Machado et al. [17] highlighted inconsistencies in behavioral outcomes, with some protocols showing no clear benefit. Holgado et al. [18] also reported no significant improvement in cycling performance following tDCS, whereas Mesquita et al. [19] observed impaired kicking performance in elite taekwondo athletes. These findings suggest that methodological heterogeneity may underlie the inconsistent effects of anodal tDCS.

One possible contributor to these inconsistent findings is that the tDCS literature includes studies with different primary aims. Some studies apply stimulation to probe causal brain–behavior relationships by expecting the stimulation to *disrupt* the performance, whereas others explicitly test performance *enhancement*. The pooling of these distinct aims may have increased heterogeneity. Therefore, the present review focuses on randomized trials that explicitly examined the enhancement effect of anodal tDCS on motor performance, which should be the case for any statistical analysis. By integrating findings from randomized controlled trials (RCTs), this meta-analysis aimed to provide an evidence-based assessment of the conditions under which anodal tDCS may improve motor performance in healthy adults, thereby informing scientifically grounded use and future translational research.

## 2. METHODS

### 2.1. Study design

This systematic review and meta-analysis were conducted in accordance with the Minds Handbook for Clinical Practice Guideline Development 2020 version 3.0, chapter 4 (Minds Manual Developing Committee, 2021). The PICO criteria for the current study were as follows: population, healthy human adults aged 18–65 years; intervention, anodal tDCS to enhance motor performance; comparators, any control condition, including sham stimulation, conventional training aimed at improving the target motor skill (e.g., physical training), or a no-intervention group; and outcomes, motor performance following anodal tDCS.

### 2.2. Database search

Systematic searches were conducted in PubMed, Scopus, Web of Science, JDreamIII, and Ichushi-Web databases to identify peer-reviewed articles published in English and Japanese from January 1, 1990 to May 25, 2022. Two authors (AS and AM) independently searched the databases. The search queries for each database are available at OSF (https://osf.io/6kghf/). In brief, the search strategy encompassed two main domains: terms related to “non-invasive brain stimulation” and terms related to “motor performance”. For “motor performance”-related terms, three sets of terms were included: (1) “improve”-related terms (e.g., “advance” and “enhance”), (2) “motor”-related terms (e.g., “sports” and “athletic”), and (3) “skill”-related terms (e.g., “accuracy” and “speed”). Within each set of terms, the terms were combined with the “OR” function on search engines. Then, the four “OR”-combined sets of terms were combined with the “AND” function. The asterisk function (e.g., “muscl*”) was used to capture all variations of the terms in the search engines. English and Japanese terms were used in the Japanese databases.

### 2.3. Study selection

Articles were screened in two phases. The first screening step involved the removal of irrelevant articles based on the relevance of the titles and abstracts. The second screening step was a full-text review to assess eligibility according to the predefined criteria described later. Search results were merged into a single library, duplicates were removed, and additional papers identified from other sources were included. Subsequently, AS, AM, and YG independently performed the first screening based on title and abstract relevance. Studies with disagreements between the individual judgments were included in the full-text assessment (second screening). AS, AM, and YG independently performed the second screening, and TI, NY, and NH conducted a full-text assessment of studies with disagreements regarding eligibility in the second screening. Four reviewers (AS, TI, NY, and NH) discussed and resolved any discrepancies until a consensus was reached.

Studies included in the qualitative synthesis met all of the following eligibility criteria: (i) participants were healthy adults aged 18–65 years; (ii) the intervention involved anodal tDCS intended to enhance motor performance; (iii) motor performance was measured as an outcome, where any observable motor output was considered; (iv) the study included a comparator condition enabling estimation of the tDCS effect; (v) the study was an original article published in a peer-reviewed scholarly journal, written in English or Japanese; (vi) the study stated an explicit hypothesis or aim regarding motor performance improvement following anodal tDCS; and (vii) the study design was a parallel-group RCT. We excluded crossover randomized trials because within-participant dependencies and potential carry-over effects complicate standardized extraction and interpretation of heterogeneous motor outcomes. We also excluded studies in which non-tDCS stimulation was applied before performance evaluation (e.g., TMS before outcome assessment).

Additionally, for studies included in the meta-analysis, the following criteria were required: (viii) sufficient outcome data were available to calculate an effect size, (ix) duplicate datasets were excluded, and (x) the study was assessed as having a low or moderate risk of bias.

### 2.4. Data extraction

AS, AM, and YG independently extracted the following data during the second screening: study design, participant demographics, types of interventions, control conditions, and outcome measures related to behavioral motor performance. For studies included in the qualitative synthesis, AS and TI further extracted detailed information about interventions and experimental protocols as well as descriptions of serious adverse events and side effects.

For the meta-analysis, motor performance measures following the intervention were extracted from the RCTs. Each eligible comparison in a study was treated as a separate trial. For each trial, motor performance data were extracted as means, standard deviations, and the number of participants in each group. The extracted data were recorded in a predesigned spreadsheet that automatically calculated the standardized mean difference (SMD; Hedges’ g, which corrects for small sample bias) as an effect size. We contacted the authors to request missing outcome data and the necessary details of the study.

### 2.5. Risk of bias assessment

AS and TI independently assessed the risk of bias in studies from which the statistics of motor performance measures were extracted. The assessment was conducted using the Cochrane Collaboration’s tool [20]. For each study, the risk of bias was categorized as low, high, or unclear across the following six domains: (i) selection bias, based on random allocation sequence, baseline imbalance, and allocation concealment; (ii) performance bias, based on blinding of participants and personnel; (iii) detection bias, based on blinding of outcome assessments; (iv) attrition bias, based on the incompleteness of outcome data and intention-to-treat analysis; (v) reporting bias, based on selective reporting; and (vi) other possible sources of bias, such as conflict of interest (COI), a sample size determination method, and incorrect statistical methods. Disagreements were resolved through discussion among the four authors (AS, TI, NY, and NH). Studies with a high risk of bias in more than two domains were excluded from the meta-analysis.

### 2.6. Data analysis

For studies that passed full-text screening, we summarized and visualized the study and sample characteristics. For studies that met the risk-of-bias criteria, we conducted a standard pairwise meta-analysis using a random-effects model by combining all data of the same outcome into a single dataset to avoid selective analysis of outcomes. Since multiple dependent effect sizes were obtained from the same study, robust variance estimation [21] was performed to deal with the dependent effect sizes. Cluster-robust variance estimation with small-sample corrections [22] was implemented using the clubSandwich package. Because this approach directly estimates cluster-robust variance, no assumption regarding the within-study correlation (ρ) among effect sizes was required. Publication bias was visualized as a funnel plot and estimated using the Begg and Egger test. To evaluate the effect of outlying/influential data points due to the small sample size, the results of meta-analyses using a random-effects model and a fixed-effects model were compared [23].

This meta-analysis included only studies that hypothesized that anodal tDCS enhances motor performance. Consequently, we used one-tailed tests to assess the statistical significance of the effects in the pairwise meta-analysis. With this clear hypothesis, the results are presented in terms of whether the stimulation condition showed a significantly greater positive effect than the control condition, accompanied by a 95% lower confidence bound (LCB) to represent the lower limit of the directional effect estimate.

Heterogeneity across the studies was assessed using Cochran Q and I^2^ statistics. We conducted subgroup meta-analyses to clarify the efficacy of anodal tDCS for the following categorical items: (i) timing of electrical stimulation; (ii) timing of motor performance assessment; (iii) type of motor performance; and (iv) location of electrical stimulation. To explore the dose-response relationship, a meta-regression analysis was applied to examine the correlation between the effect size and the intensity of anodal tDCS and the correlation between the effect size and the product of total stimulation duration and intensity in each study. We defined the total stimulation duration as the duration of each session × the number of sessions. If the stimulation duration in each session was described as a range, the mid-value was used.

All statistical analyses were performed using metafor (version 3.4.0), meta (version 5.5.0), clubSandwich (version 0.5.8), and dmetar (version 0.1.0) in R (version 4.2.1; https://www.r-project.org/). Meta-analysis, subgroup analysis, meta-regression analysis, and related visualizations were conducted using the metafor package in R (version 4.2.1). Robust variance estimation was performed using the robust function with the metafor and clubSandwich packages in R (version 4.2.1).

## 3. RESULTS

### 3.1. Search results

We obtained 3,796 records through a database search and 5 additional records from the reference sections of the retrieved studies. After title/abstract screening, 3,448 records were excluded. Full texts of the remaining 353 reports were assessed for eligibility. Of these, 328 reports were excluded for the following reasons: non-original article (n=14); inclusion of patient populations (n=3); participants outside the target age range (n=6); absence of motor performance outcomes (n=37); lack of a hypothesis regarding motor improvement (n=97); non-RCT design (n=26); crossover design (n=105); use of non-tDCS stimulation (n=11); incomplete data (n=28), and duplicate data (n=1).

Accordingly, 25 studies were included in the qualitative synthesis. Two studies were excluded from the meta-analysis because of a high risk of bias, yielding 23 studies that contributed 25 trials (N=885) to the meta-analysis. The overall study selection process is shown in Figure 1.

**Fig. 1.**
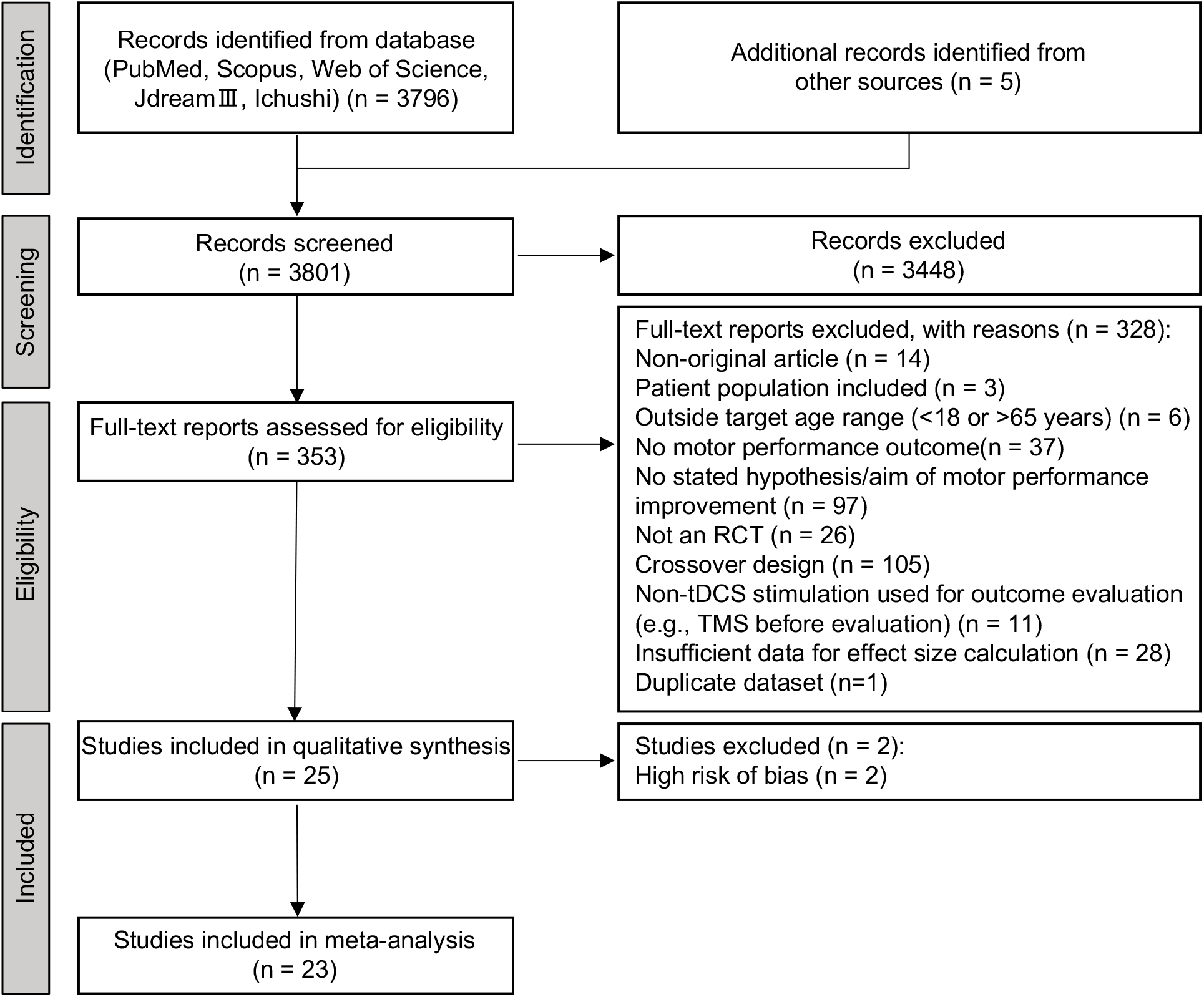
Preferred Reporting Items for Systematic Reviews and Meta-Analyses flowchart diagram of the literature search and study selection.

### 3.2. Study and sample characteristics

Table 1 summarizes the main characteristics of all the studies included in the qualitative synthesis. The most frequently targeted sites (Fig. 2A) were the motor cortex regions in 18 studies, followed by the cerebellar regions (8 studies), frontal areas (7 studies), parietal regions (2 studies), and visual cortex (2 studies), while some studies targeted more than one site. The most common stimulation intensity (Fig. 2C) was 2 mA (14 studies), followed by 1 mA (7 studies) and 1.5 mA (6 studies).

**Table 1.** Characteristics of studies included in the qualitative synthesis.

| Study | Participants | Target site (anode) | Duration/day | Intervention period | Timing | Intensity | Effector | Task |
| --- | --- | --- | --- | --- | --- | --- | --- | --- |
| Apšvalka et al., 2018 [31] | Right-handed university students | M1 (C4) | 20 min | 4 days | During training | 1 mA | Upper limb | Keypress sequence learning (sequential key pressing) |
| Ciechanski et al., 2017 [32] | Medical students | M1 (C3) | 20 min | 1 day | During training | 1 mA | Upper limb | Surgical task |
| Ciechanski et al., 2019 [33] | Healthy adults (medical and veterinary students) | CP3 or CP4 (dominant hemisphere) | 20 min | 1 day | During training | 1 mA | Upper limb | Surgical task (Fundamentals of Laparoscopic Surgery (FLS) pattern cutting and peg transfer tasks) |
| Cox et al., 2020 [34] | Healthy adults | M1 (C3; left of vertex) / SMA (Cz; anterior to vertex) | 20 min x 2 | 3 days | During training | 2 mA | Upper limb | Surgical task (Fundamentals of Laparoscopic Surgery (FLS) Peg transfer task) |
| Doppelmayr et al., 2016 [35] | Healthy adults | Cerebellum (midline; below Oz) / Parietal (P4) / M1 (C3) | 21 min | 1 day | During training | 1 mA | Upper limb | Motor adaptation during mirror drawing tasks |
| Ehsani et al., 2016 [36] | Healthy university students | M1 (C3) / Cerebellum (ipsilateral; 1 cm below the inion, 1 cm medial to the mastoid) | 20 min | 1 day | During training | 2 mA | Upper limb | Serial reaction time task (SRTT) |
| Harris et al., 2019 [37] | Healthy adults (all right-handed novice golfers) | F4 / C4 / Oz | 5 min pre-task + during task performance | 1 day | Pre-task (5 min) + during training | 1.5 mA | Whole body | Golf putting |
| Jackson et al., 2019 [38] | Healthy adult males (no recent throwing sports) | 3 cm right of the inion | 25 min | 1 day | During training | 2 mA | Upper limb | Overhand throwing task |
| Kico et al., 2022 [39] | Healthy adults | M1 (Cz) | 20 min | 1 day | Before training | 1.5 mA | Whole body | Dance |
| Kumari et al., 2020 [40] | Healthy adults | Cerebellum (3 cm lateral to the inion; ipsilateral to dominant leg) | 15 min | 3 days | During task performance | 2 mA | Whole body | Split belt treadmill walking |
| Labruna et al., 2019 [41] | Healthy adults | M1 | 20 min | 1 day | During training | 2 mA | Upper limb | Visuomotor adaptation task |
| Lerner et al., 2021 [42] | Right-handed young healthy adults | C4, F1 | 20 min | 1 day | During training | 2 mA<br>1.5 mA | Upper limb | Motor sequence learning task |
| Liew et al., 2018 [43] | Healthy adults | Cerebellum (right; 3 cm lateral to inion) / PFC (F3) | 25 min | 1 day | During task performance | 2 mA | Upper limb | Visuomotor adaptation task |
| Lum et al., 2018 [44] | Right-handed healthy adults | centered over the intersection between T3-Fz and F7-Cz | 16 min | 1 day | During training | 2 mA | Upper limb | Serial reaction time task (SRTT) |
| Maeda et al., 2017 [45] | Not specified | M1(TMS hotspot; non-dominant leg representation) | 10 min | 7 days | During training | 2 mA | Lower limb | Strength training |
| Molero-Chamizo et al., 2018 [46] | Right-handed adults | M1 (C3) | 15 min | 1 day | Before task performance | 1.5 mA | Upper limb | Go/no-go response task |
| Neto et al., 2022 [47] | Amateur soccer players | F3 | 20 min | 5 days | During training | 2 mA | Whole body | Choice reaction time task |
| Nguemeni et al., 2021 [48] | Healthy adults | Cerebellum (left cerebellar cortex; 3 cm lateral to theinion) | 20 min | 1 day | Before / during / after training | 2 mA | Upper limb | Sequential tapping task (sequential key pressing) |
| Orcioli-Silva et al., 2021 [49] | Healthy adults | M1 (Cz) / IPFC (AF3-FP1) | 20 min | 1 day | During training | 0.6 mA | Whole body | Treadmill walking |
| Parma et al., 2021 [50] | Healthy adults | M1 (C3) | 20 min | 2 days | Before training | 1.5 mA | Whole body | Golf putting |
| Saimpont et al., 2016 [51] | Right-handed healthy adults | M1 (C4) | 13 min | 1 day | During training | 2 mA | Upper limb | Finger tapping sequence |
| Salvador et al., 2017 [52] | Right-handed university students | M1 (C4) | 20 min | 1 day | Before task performance | 1 mA | Upper limb | Pegboard task |
| Sevilla-Sanchez et al., 2021 [53] | Healthy adults | M1 (C3) | 15 min | 20 days | During training | 1.5 mA | Upper limb | Typing |
| Talimkhani et al., 2019 [54] | Healthy adults | Dual-site anodal tDCS (left DLPFC + M1); control: M1 | 20 min | 3 days | During training | 1 mA | Upper limb | Serial reaction time task (SRTT) |
| von Rein et al., 2015 [55] | Healthy adults | M1 (right hemisphere) | 20 min | 1 day | During training | 1 mA | Upper limb | Ball rotation task |

M1, primary motor cortex; SMA, supplementary motor area; DLPFC, dorsolateral prefrontal cortex; IPFC, left prefrontal cortex.

**Fig. 2.**
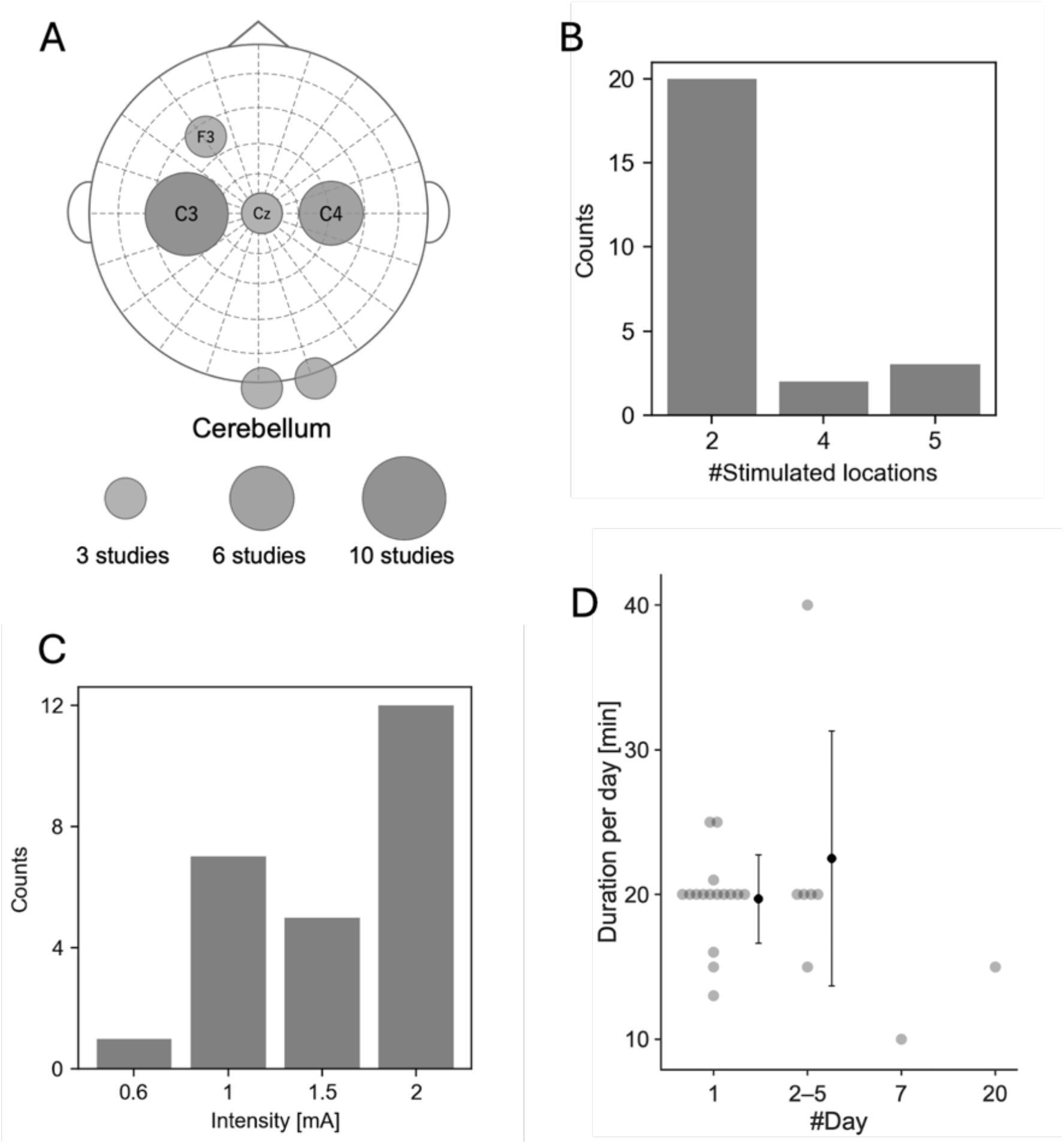
Qualitative summary of the included studies. (A) Distribution of stimulation targets, where the size of each node represents the number of studies. (B) Number of stimulated locations. (C) Stimulation intensity. (D) Daily stimulation duration grouped by the length of the intervention period. Gray dots indicate individual trials. Black dots and error bars represent the mean and the standard deviation of each group.

Most studies targeted upper-limb tasks (18 studies), 6 studies examined whole-body tasks, and 1 study focused on lower-limb tasks. Common task paradigms included reaction-time tasks (six studies), motor adaptation tasks such as visuomotor adaptation (five studies), sports-related tasks such as golf putting and dance (three studies), and surgical tasks (three studies).

Most studies applied stimulation for a single day (19 studies), 3 studies used 3-day protocols, and 1 study used 2-, 4-, 5-, 7-, or 20-day protocols (Fig. 2D). Stimulation was most commonly applied during task practice/training (20 studies), followed by during task performance/assessment (4 studies) and before task performance/assessment (2 studies), with several studies using other timing protocols.

### 3.3. Risk-of-bias assessment

The risk-of-bias assessment for the 25 included studies is summarized in Figure 3. Two studies judged to have a high risk of bias in more than two domains were excluded from the meta-analysis.

**Fig. 3.**
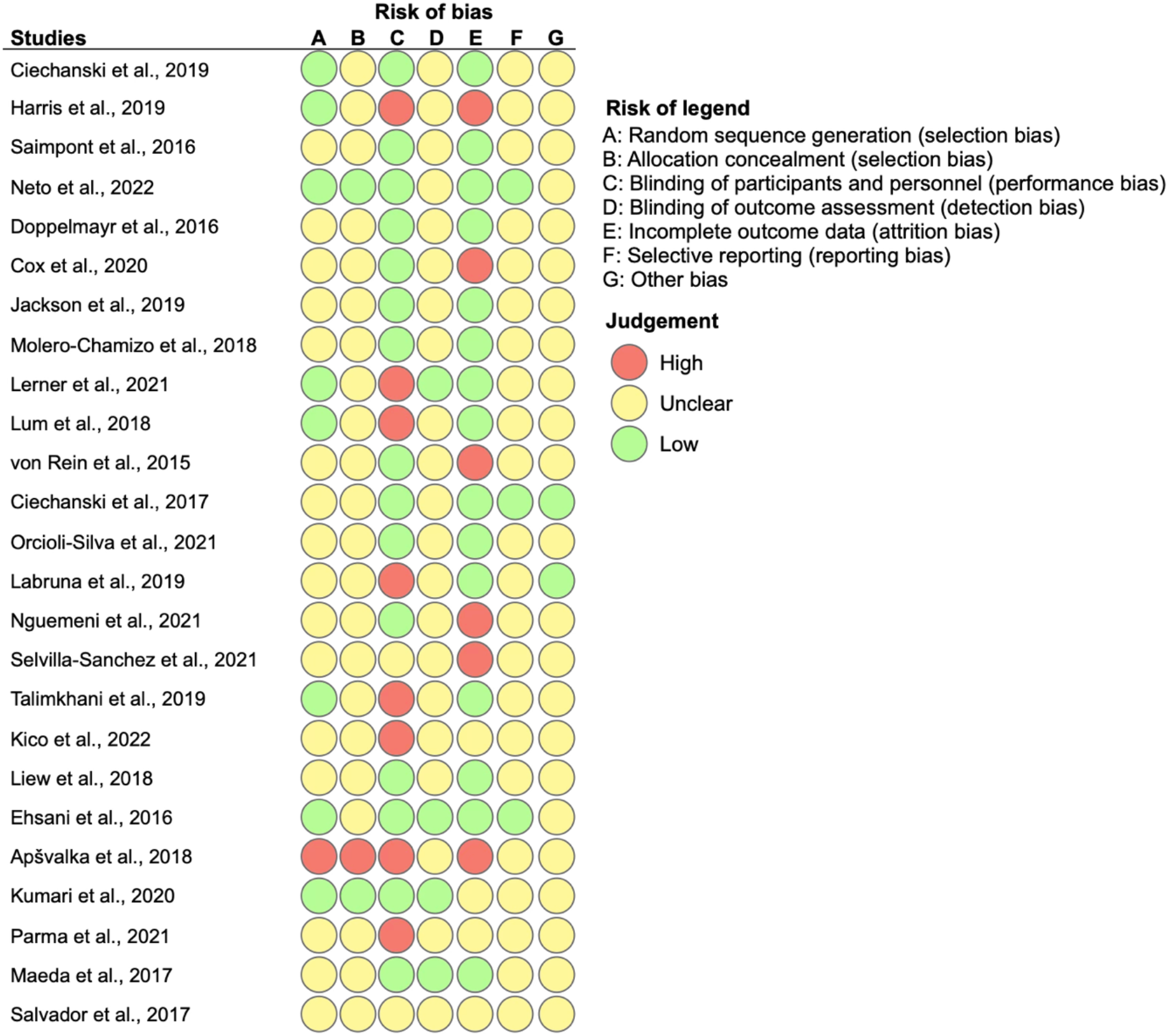
Risk of bias across all included studies. Judgments are shown for each domain (A-G). Green, yellow, and red indicate low, unclear, and high risk of bias, respectively.

### 3.4. Overall effects of anodal tDCS for improving motor performance

Twenty-five trials (N=885) from 23 studies were included in the meta-analysis using a random-effects model. Overall, anodal tDCS had a moderate effect on motor performance (SMD=0.67, 95% LCB=0.31, t=3.35, P=0.003; Fig. 4). However, caution is needed when interpreting the results, since there was significant asymmetry in the funnel plot (Fig. 5; Begg’s test: tau=0.38, p<.0001; Egger’s test: z=7.35, p<.001), indicating the existence of publication bias. There was significant heterogeneity across the trials (Q=404.3, P<0.001, I^2^=80.0%).

**Fig. 4.**
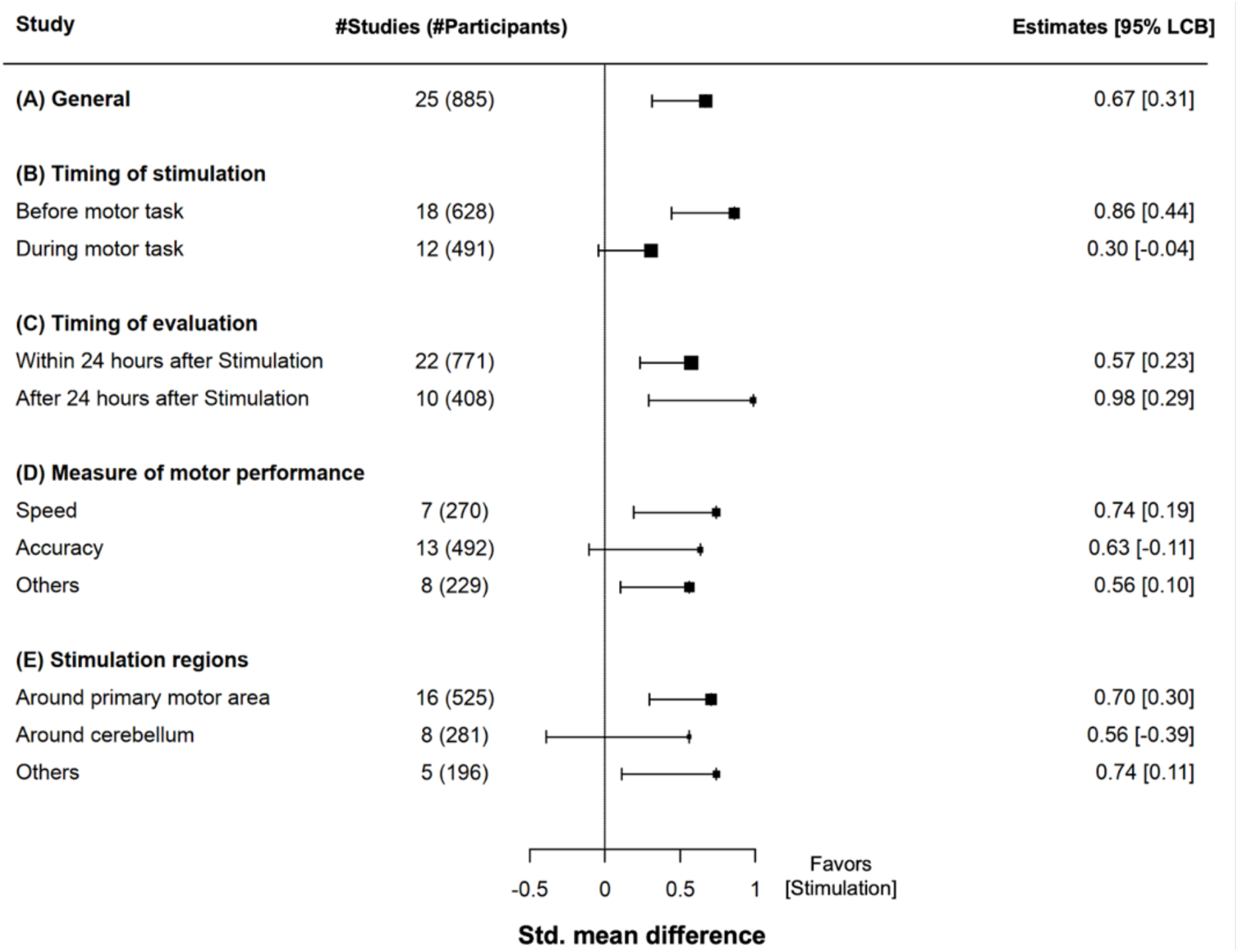
Forest plot of the effects of anodal transcranial direct current stimulation on motor performance. Squares and horizontal lines indicate the standardized mean difference and its 95% lower confidence bound, respectively, consistent with the one-tailed framework described in methods. (A) Overall effect size estimate. (B-E) Effect size estimate from subgroup analyses.

**Fig. 5.**
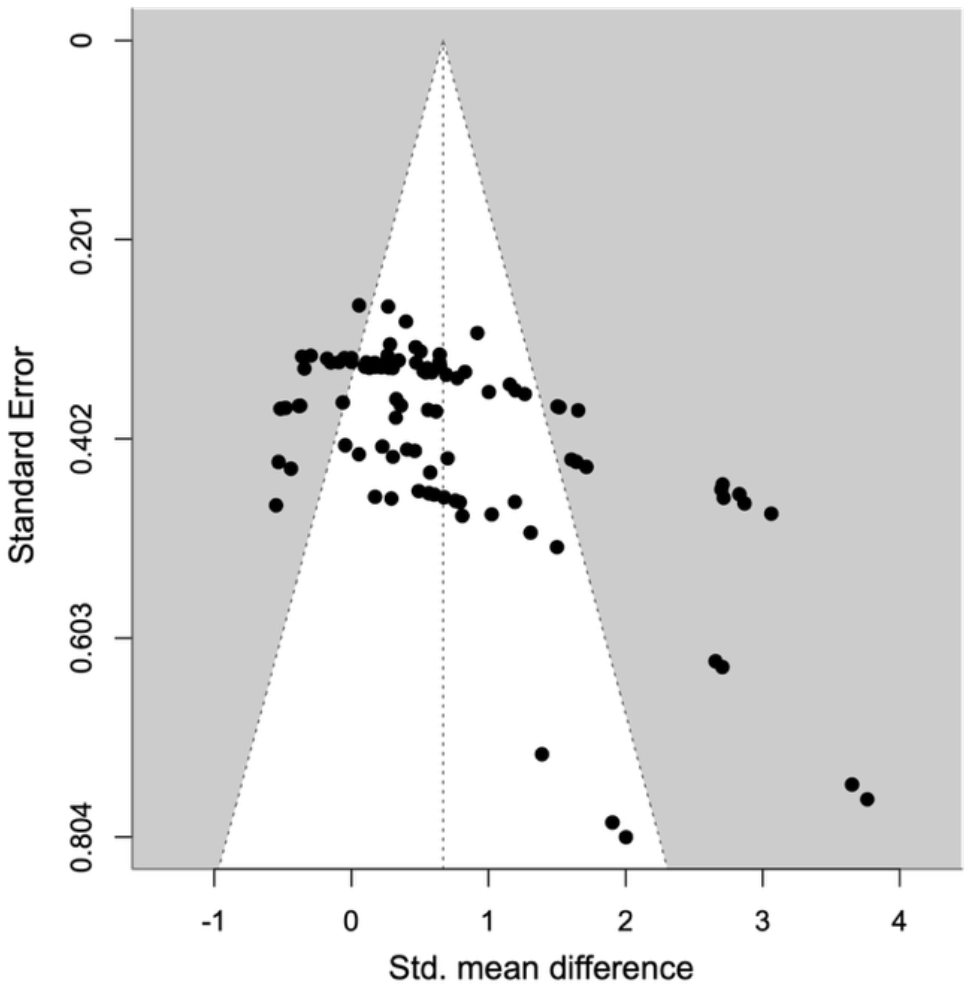
Funnel plot displaying effect sizes from the experimental results.

For sensitivity analysis, we compared the results of the meta-analyses using random-effects and fixed-effects models (SMD=0.54, 95% LCB=0.24, t=3.24, P=0.004) to evaluate whether the effect was contaminated by outlying/influential data points. The results of both analyses were comparable, indicating the robustness of the result across different analyses.

### 3.5.Subgroup meta-analysis

We performed subgroup meta-analyses to evaluate the effect of anodal tDCS for the following categorical items: (i) timing of electrical stimulation (before or during motor task); (ii) timing of motor performance assessment (within 24 hours post-stimulation or beyond 24 hours post-stimulation); (iii) type of motor performance (speed, accuracy, or others); (iv) location of electrical stimulation (around primary motor area, around cerebellum, or others). Because some studies contributed to multiple trials, subgroup trial counts can exceed the total number of unique trials in the primary analysis; thus, statistical dependency was handled using robust variance estimation.

#### 3.5.1. Effects of stimulation timing

The estimated effect of stimulation timing before a motor task (18 trials) was large (SMD=0.86, 95% LCB=0.44, t=3.74, P=0.004; heterogeneity: Q=286.8, P<.001, I^2^=80.9%), and that during a motor task (12 trials) was not significant (SMD=0.30, 95% LCB=–0.04, t=1.67, P=0.07; heterogeneity: Q=92.5, P< 0.001, I^2^=69.3%).

#### 3.5.2. Effects of timing of motor performance evaluation after stimulation

The estimated effect within 24 hours post-stimulation (22 trials) was moderate (SMD=0.57, 95% LCB=0.23, t=3.08, P=0.006; heterogeneity: Q=247.7, P<0.001, I^2^=73.4%). By contrast, the estimated effect beyond 24 hours post-stimulation (10 trials) was large (SMD=0.98, 95% LCB=0.29, t=2.76, P=0.017; heterogeneity: Q=148.7, P<0.001, I^2^=88.7%).

#### 3.5.3. Effects of differences in the motor performance measures

The estimated effect size for speed (7 trials) was medium (SMD=0.74, 95% LCB=0.19, t=2.89, P=0.023; heterogeneity: Q=132.0, P<0.0001, I^2^=70.1%). For accuracy (13 trials), the effect size was not significant (SMD=0.63, 95% LCB=–0.11, t=1.61, P=0.150; heterogeneity: Q=201.7, P<0.001, I^2^=85.6%). Moreover, the “others” category (e.g., composite scores or qualitative measures; 8 trials) showed a medium effect (SMD=0.56, 95% LCB=0.10, t=2.40, P=0.028; heterogeneity: Q=64.5, P<0.001, I^2^=81.0%).

#### 3.5.4. Effects of differences in the stimulation regions

The subgroup stimulated around M1 (16 trials) demonstrated a medium effect size (SMD=0.70, 95% LCB=0.30, t=3.23, P=0.065; heterogeneity: Q=220.8, P<0.001, I^2^=77.7%), whereas cerebellar stimulation (8 trials) yielded no significant effect size (SMD=0.56, 95% LCB=–0.39, t=1.25, P=0.140; heterogeneity: Q=151.2, P<0.001, I^2^=87.7%). In the “other” regions subgroup (5 trials), the effect was medium (SMD=0.74, 95% LCB=0.11, t=2.84, P=0.035; heterogeneity: Q=28.9, P=0.002, I^2^=62.9%).

### 3.6. Meta-regression analysis

Meta-regression analysis revealed no significant relationship between effect size and the intensity of anodal tDCS (25 trials; beta=0.1452, 95% CI=−0.7436–1.0339, t=0.3918, P=0.71) or the product of total stimulation duration and intensity (25 trials; beta=−0.0016, 95% CI= −0.0079–0.0047, t=−0.8329, P=0.47). The meta-regression analysis for all other subgroups yielded no significant results (all P>0.1).

### 3.7. Adverse effects

Adverse events related to anodal tDCS were assessed in 12 trials using scale indicators, such as the visual analog or numerical rating scale, and 4 trials reported the number of cases of adverse events. None of the trials reported serious adverse events.

Analysis of scale indicators revealed that the tingling/itching sensation was not significantly different between the tDCS and control groups (Tingling: 5 trials, SMD=0.97, 95% CI [–0.28, 2.21], t=2.29, P=0.094; heterogeneity: Q=132.0, P<0.0001, I^2^=93.3%; Itching: 4 trials, SMD=1.23, 95% CI [– 1.92, 4.38], t=1.71, P=0.23; heterogeneity: Q=87.4, P<0.0001, I^2^=93.8%).

## 4. DISCUSSION

This meta-analysis evaluated the efficacy of anodal tDCS for improving motor performance in healthy adults. While our results suggest a positive average effect of anodal tDCS on certain motor outcomes, they also revealed substantial heterogeneity and methodological limitations across studies, suggesting caution in interpreting the findings.

Several studies included in this review reported performance enhancements in strength, endurance, and motor learning following anodal tDCS. Importantly, this review focused on studies that explicitly hypothesized the performance-enhancing effect of anodal tDCS. This criterion is critical in distinguishing studies designed to evaluate improvement from those assessing other effects, such as interference or neutral outcomes. During the screening process, many studies were excluded because they lacked this explicit hypothesis, thereby reducing the risk of conflating heterogeneous objectives and ensuring a more focused and interpretable synthesis. However, the reliability of these findings remains unclear. Many tDCS studies still lack essential controls such as proper randomization, blinding, and adequate power calculations. Funnel plot asymmetry and high inter-study heterogeneity further raise concerns about potential publication bias and variability in the experimental design. Taken together, these findings suggest that the literature is in an early or exploratory phase. More rigorously designed and adequately powered RCTs are essential to establish the generalizability and robustness of the effects of anodal tDCS on motor performance.

Mechanistically, the primary physiological effect of anodal tDCS is subthreshold depolarization of the neuronal membrane, which increases cortical excitability. Nitsche and Paulus [8] demonstrated that direct current stimulation of the motor cortex can modulate excitability as indicated by increased motor-evoked potential (MEP) amplitudes. Anodal tDCS typically facilitates excitability, whereas cathodal tDCS induces suppression. These changes are thought to reflect shifts in the excitatory/inhibitory balance of cortical circuits [10], potentially leading to synaptic modifications resembling long-term potentiation (LTP). However, evidence suggests that the magnitude and direction of MEP modulation are not linearly related to stimulation intensity or duration [24–26]. For instance, Jamil et al. [25] systematically tested stimulation intensities from 0.5 to 2.0 mA and found that anodal tDCS-induced MEP facilitation did not increase linearly with stimulation intensity, whereas Hsu et al. [26] observed no monotonic gain in excitability at stimulation intensities up to 6 mA. These findings challenge the assumption of a linear dose-response relationship at the physiological level. Consistent with these findings, our meta-regression analyses revealed no significant association between stimulation intensity or cumulative dose and behavioral effect size. However, the available dose range was narrow, and the heterogeneity was high. Therefore, these null findings should be interpreted as inconclusive rather than as evidence against a dose-response relationship. This suggests a potential decoupling between physiological and behavioral outcomes: MEP modulation may not reliably predict motor performance gains, and the behavioral efficacy of tDCS likely depends on additional factors such as task characteristics, stimulation timing, and individual neurophysiological states. Although MEPs offer valuable insights into cortical excitability [8], they may be insufficient as stand-alone predictors of functional outcomes. Elucidating the mechanisms by which tDCS-induced excitability changes translate into behavioral improvements remains a key challenge in the field. Thus, increases in MEP amplitude after anodal tDCS should not be interpreted as reliable biomarkers of improved motor performance in healthy adults.

We conducted subgroup analyses to explore the potential factors influencing the effectiveness of tDCS. Although no single factor consistently explained the observed effects across all studies, several patterns emerged. For instance, stimulation delivered before motor task execution tended to produce greater effects than stimulation delivered during the task. This finding aligns with that of Simonsmeier et al.’s study [27], who reported that stimulation during the learning phase, rather than the test phase, was more effective in enhancing cognitive outcomes. It is plausible that in motor tasks, pre-task stimulation facilitates neuroplastic changes that enhance subsequent execution. Additionally, motor performance assessed beyond 24 hours post-stimulation showed greater effects than those measured within 24 hours. This delayed benefit may reflect consolidation processes, consistent with the findings of Hashemirad et al. [15], who observed more pronounced retention phase improvements in motor sequence learning following anodal tDCS. These results suggest that tDCS may not act immediately but rather support longer-term changes in neural function. Task characteristics also influenced the outcomes. Tasks involving speed-based or reactive movements tended to show more pronounced improvements than those requiring precise motor control. These findings suggest that the type of motor task plays a key role in determining the extent to which anodal tDCS can modulate performance. This interpretation is broadly consistent with prior work by Maudrich et al. [16], who reported that tDCS effects varied across motor domains, with significant effects in visuomotor tasks but not in strength- or endurance-based measures. Finally, stimulation over the M1 was associated with more consistent improvements in motor performance than stimulation over the cerebellum or other areas. Nevertheless, even among M1-targeted studies, the variability in outcomes was substantial.

These findings suggest that the effectiveness of anodal tDCS is influenced by a combination of factors, including when the stimulation is applied, which brain area is targeted, what kind of motor task is involved, and when performance is measured. However, the robustness of these effects was limited, and our meta-regression failed to identify reliable dose-related moderators. As such, while the patterns observed here and in prior systematic reviews indicate promising directions for improving stimulation strategies, the current evidence base is not sufficiently strong to guide definitive protocols. In other words, increasing stimulation intensity or cumulative dose does not guarantee additional behavioral benefit. Therefore, tDCS should be used with caution, particularly until the optimal parameters and individual response factors are better understood.

In conclusion, while anodal tDCS holds promise as a tool for improving motor performance, the present findings, together with previous evidence, suggest that its effects remain inconsistent and uncertain. Future research should prioritize individualized state-dependent stimulation approaches that consider the neurophysiological profile of each participant. Beyond traditional protocols, emerging techniques such as high-definition tDCS [28], transcranial alternating current stimulation [29], and temporal interference stimulation [30] offer new opportunities to refine spatial and temporal targeting. Comparative studies examining these modalities are essential for determining the optimal strategies for motor enhancement.

## ACKNOWLEDGEMENTS

We thank the members of the Evidence Evaluation Committee for Braintech Guidebook, a specially organized group for the Moonshot R&D project, for their comments on this manuscript.

## AUTHOR CONTRIBUTIONS

**Atsushi Sasaki**: Conceptualization, Data curation, Investigation, Methodology, Project administration, Software, Validation, Visualization, Writing – original draft, Writing – review and editing.

**Takuya Ideriha:** Conceptualization, Data curation, Formal analysis, Investigation, Methodology, Project administration, Software, Validation, Visualization, Writing – original draft, Writing – review and editing.

**Atsuya Matsuoka:** Data curation, Investigation, Validation, Writing – review and editing.

**Yujin Goto:** Data curation, Investigation, Validation, Writing – review and editing.

**Natsue Yoshimura:** Conceptualization, Data curation, Formal analysis, Investigation, Methodology, Supervision, Validation, Visualization, Writing – original draft, Writing – review and editing.

**Nobuhiro Hagura:** Conceptualization, Data curation, Formal analysis, Investigation, Methodology, Project administration, Software, Supervision, Validation, Visualization, Writing – original draft, Writing – review and editing.

## DATA STATEMENT

Data and codes used in this study were shared in OSF (https://osf.io/6kghf/).

## REFERENCES

[1] Polanía R, Nitsche MA, Ruff CC. Studying and modifying brain function with non-invasive brain stimulation. Nat Neurosci 2018;21:174–87. 10.1038/s41593-017-0054-4.

[2] Bolognini N, Pascual-Leone A, Fregni F. Using non-invasive brain stimulation to augment motor training-induced plasticity. J NeuroEngineering Rehabil 2009;6:8. 10.1186/1743-0003-6-8.

[3] Hummel FC, Cohen LG. Non-invasive brain stimulation: a new strategy to improve neurorehabilitation after stroke? The Lancet Neurology 2006;5:708–12. 10.1016/S1474-4422(06)70525-7.

[4] Reardon S. “Brain doping” may improve athletes’ performance. Nature 2016;531:283–4. 10.1038/nature.2016.19534.

[5] Elahi B, Elahi B, Chen R. Effect of transcranial magnetic stimulation on Parkinson motor function—Systematic review of controlled clinical trials. Movement Disorders 2009;24:357–63. 10.1002/mds.22364.

[6] Hsu W-Y, Cheng C-H, Liao K-K, Lee I-H, Lin Y-Y. Effects of Repetitive Transcranial Magnetic Stimulation on Motor Functions in Patients With Stroke. Stroke 2012;43:1849–57. 10.1161/STROKEAHA.111.649756.

[7] Fitzgerald PB, Fountain S, Daskalakis ZJ. A comprehensive review of the effects of rTMS on motor cortical excitability and inhibition. Clinical Neurophysiology 2006;117:2584–96. 10.1016/j.clinph.2006.06.712.

[8] Nitsche MA, Paulus W. Excitability changes induced in the human motor cortex by weak transcranial direct current stimulation. The Journal of Physiology 2000;527:633–9. 10.1111/j.1469-7793.2000.t01-1-00633.x.

[9] Dubljević V, Saigle V, Racine E. The Rising Tide of tDCS in the Media and Academic Literature. Neuron 2014;82:731–6. 10.1016/j.neuron.2014.05.003.

[10] Lefaucheur J-P, Antal A, Ayache SS, Benninger DH, Brunelin J, Cogiamanian F, et al. Evidence-based guidelines on the therapeutic use of transcranial direct current stimulation (tDCS). Clinical Neurophysiology 2017;128:56–92. 10.1016/j.clinph.2016.10.087.

[11] Pascual-Leone A. Transcranial magnetic stimulation in cognitive neuroscience – virtual lesion, chronometry, and functional connectivity. Current Opinion in Neurobiology 2000;10:232–7. 10.1016/S0959-4388(00)00081-7.

[12] Parkin BL, Ekhtiari H, Walsh VF. Non-invasive Human Brain Stimulation in Cognitive Neuroscience: A Primer. Neuron 2015;87:932–45. 10.1016/j.neuron.2015.07.032.

[13] Horvath JC, Forte JD, Carter O. Evidence that transcranial direct current stimulation (tDCS) generates little-to-no reliable neurophysiologic effect beyond MEP amplitude modulation in healthy human subjects: A systematic review. Neuropsychologia 2015;66:213–36. 10.1016/j.neuropsychologia.2014.11.021.

[14] Bikson M, Grossman P, Thomas C, Zannou AL, Jiang J, Adnan T, et al. Safety of Transcranial Direct Current Stimulation: Evidence Based Update 2016. Brain Stimulation 2016;9:641–61. 10.1016/j.brs.2016.06.004.

[15] Hashemirad F, Zoghi M, Fitzgerald PB, Jaberzadeh S. The effect of anodal transcranial direct current stimulation on motor sequence learning in healthy individuals: A systematic review and meta-analysis. Brain Cogn 2016;102:1–12. 10.1016/j.bandc.2015.11.005.

[16] Maudrich T, Ragert P, Perrey S, Kenville R. Single-session anodal transcranial direct current stimulation to enhance sport-specific performance in athletes: A systematic review and meta-analysis. Brain Stimulation 2022;15:1517–29. 10.1016/j.brs.2022.11.007.

[17] Machado DG da S, Unal G, Andrade SM, Moreira A, Altimari LR, Brunoni AR, et al. Effect of transcranial direct current stimulation on exercise performance: A systematic review and meta-analysis. Brain Stimul 2019;12:593–605. 10.1016/j.brs.2018.12.227.

[18] Holgado D, Zandonai T, Ciria LF, Zabala M, Hopker J, Sanabria D. Transcranial direct current stimulation (tDCS) over the left prefrontal cortex does not affect time-trial self-paced cycling performance: Evidence from oscillatory brain activity and power output. PLoS ONE 2019;14:e0210873. 10.1371/journal.pone.0210873.

[19] Mesquita PHC, Lage GM, Franchini E, Romano-Silva MA, Albuquerque MR. Bi-hemispheric anodal transcranial direct current stimulation worsens taekwondo-related performance. Hum Mov Sci 2019;66:578–86. 10.1016/j.humov.2019.06.003.

[20] Higgins JP, Altman DG, Gøtzsche PC, Jüni P, Moher D, Oxman AD, et al. The Cochrane Collaboration’s tool for assessing risk of bias in randomised trials. Bmj 2011;343.

[21] Hedges LV, Tipton E, Johnson MC. Robust variance estimation in meta-regression with dependent effect size estimates. Research Synthesis Methods 2010;1:39–65. 10.1002/jrsm.5.

[22] Tipton E. Small sample adjustments for robust variance estimation with meta-regression. Psychological Methods 2015;20:375–93. 10.1037/met0000011.

[23] Higgins JP, Green S. Cochrane handbook for systematic reviews of interventions 2008.

[24] Batsikadze G, Moliadze V, Paulus W, Kuo M-F, Nitsche MA. Partially non-linear stimulation intensity-dependent effects of direct current stimulation on motor cortex excitability in humans. J Physiol 2013;591:1987–2000. 10.1113/jphysiol.2012.249730.

[25] Jamil A, Batsikadze G, Kuo H-I, Labruna L, Hasan A, Paulus W, et al. Systematic evaluation of the impact of stimulation intensity on neuroplastic after-effects induced by transcranial direct current stimulation. J Physiol 2017;595:1273–88. 10.1113/JP272738.

[26] Hsu G, Jafari ZH, Ahmed A, Edwards DJ, Cohen LG, Parra LC. Dose–response of tDCS effects on motor learning and cortical excitability: A preregistered study. Imaging Neuroscience 2025;3:imag_a_00431. 10.1162/imag_a_00431.

[27] Simonsmeier BA, Grabner RH, Hein J, Krenz U, Schneider M. Electrical brain stimulation (tES) improves learning more than performance: A meta-analysis. Neuroscience & Biobehavioral Reviews 2018;84:171–81. 10.1016/j.neubiorev.2017.11.001.

[28] Kuo H-I, Bikson M, Datta A, Minhas P, Paulus W, Kuo M-F, et al. Comparing Cortical Plasticity Induced by Conventional and High-Definition 4 × 1 Ring tDCS: A Neurophysiological Study. Brain Stimulation 2013;6:644–8. 10.1016/j.brs.2012.09.010.

[29] Antal A, Paulus W. Transcranial alternating current stimulation (tACS). Front Hum Neurosci 2013;7:317. 10.3389/fnhum.2013.00317.

[30] Violante IR, Alania K, Cassarà AM, Neufeld E, Acerbo E, Carron R, et al. Non-invasive temporal interference electrical stimulation of the human hippocampus. Nat Neurosci 2023;26:1994–2004. 10.1038/s41593-023-01456-8.

[31] Apšvalka D, Ramsey R, Cross ES. Anodal tDCS over Primary Motor Cortex Provides No Advantage to Learning Motor Sequences via Observation. Neural Plast 2018;2018:1237962. 10.1155/2018/1237962.

[32] Ciechanski P, Cheng A, Lopushinsky S, Hecker K, Gan LS, Lang S, et al. Effects of Transcranial Direct-Current Stimulation on Neurosurgical Skill Acquisition: A Randomized Controlled Trial. World Neurosurg 2017;108:876–884.e4. 10.1016/j.wneu.2017.08.123.

[33] Ciechanski P, Kirton A, Wilson B, Williams CC, Anderson SJ, Cheng A, et al. Electroencephalography correlates of transcranial direct-current stimulation enhanced surgical skill learning: A replication and extension study. Brain Res 2019;1725:146445. 10.1016/j.brainres.2019.146445.

[34] Cox ML, Deng Z-D, Palmer H, Watts A, Beynel L, Young JR, et al. Utilizing transcranial direct current stimulation to enhance laparoscopic technical skills training: A randomized controlled trial. Brain Stimul 2020;13:863–72. 10.1016/j.brs.2020.03.009.

[35] Doppelmayr M, Pixa NH, Steinberg F. Cerebellar, but not Motor or Parietal, High-Density Anodal Transcranial Direct Current Stimulation Facilitates Motor Adaptation. J Int Neuropsychol Soc 2016;22:928–36. 10.1017/S1355617716000345.

[36] Ehsani F, Bakhtiary AH, Jaberzadeh S, Talimkhani A, Hajihasani A. Differential effects of primary motor cortex and cerebellar transcranial direct current stimulation on motor learning in healthy individuals: A randomized double-blind sham-controlled study. Neurosci Res 2016;112:10–9. 10.1016/j.neures.2016.06.003.

[37] Harris DJ, Wilson MR, Buckingham G, Vine SJ. No effect of transcranial direct current stimulation of frontal, motor or visual cortex on performance of a self-paced visuomotor skill. Psychology of Sport and Exercise 2019;43:368–73. 10.1016/j.psychsport.2019.04.014.

[38] Jackson AK, de Albuquerque LL, Pantovic M, Fischer KM, Guadagnoli MA, Riley ZA, et al. Cerebellar Transcranial Direct Current Stimulation Enhances Motor Learning in a Complex Overhand Throwing Task. Cerebellum 2019;18:813–6. 10.1007/s12311-019-01040-6.

[39] Kico I, Liarokapis F. Enhancing the learning process of folk dances using augmented reality and non-invasive brain stimulation. Entertainment Computing 2022;40:100455. 10.1016/j.entcom.2021.100455.

[40] Kumari N, Taylor D, Rashid U, Vandal AC, Smith PF, Signal N. Cerebellar transcranial direct current stimulation for learning a novel split-belt treadmill task: a randomised controlled trial. Sci Rep 2020;10:11853. 10.1038/s41598-020-68825-2.

[41] Labruna L, Stark-Inbar A, Breska A, Dabit M, Vanderschelden B, Nitsche MA, et al. Individual differences in TMS sensitivity influence the efficacy of tDCS in facilitating sensorimotor adaptation. Brain Stimul 2019;12:992–1000. 10.1016/j.brs.2019.03.008.

[42] Lerner O, Friedman J, Frenkel-Toledo S. The effect of high-definition transcranial direct current stimulation intensity on motor performance in healthy adults: a randomized controlled trial. J Neuroeng Rehabil 2021;18:103. 10.1186/s12984-021-00899-z.

[43] Liew S-L, Thompson T, Ramirez J, Butcher PA, Taylor JA, Celnik PA. Variable Neural Contributions to Explicit and Implicit Learning During Visuomotor Adaptation. Front Neurosci 2018;12:610. 10.3389/fnins.2018.00610.

[44] Lum JAG, Mills A, Plumridge JMA, Sloan NP, Clark GM, Hedenius M, et al. Transcranial direct current stimulation enhances retention of a second (but not first) order conditional visuo-motor sequence. Brain Cogn 2018;127:34–41. 10.1016/j.bandc.2018.09.006.

[45] Maeda K, Yamaguchi T, Tatemoto T, Kondo K, Otaka Y, Tanaka S. Transcranial Direct Current Stimulation Does Not Affect Lower Extremity Muscle Strength Training in Healthy Individuals: A Triple-Blind, Sham-Controlled Study. Front Neurosci 2017;11:179. 10.3389/fnins.2017.00179.

[46] Molero-Chamizo A, Alameda Bailén JR, Garrido Béjar T, García López M, Jaén Rodríguez I, Gutiérrez Lérida C, et al. Poststimulation time interval-dependent effects of motor cortex anodal tDCS on reaction-time task performance. Cogn Affect Behav Neurosci 2018;18:167–75. 10.3758/s13415-018-0561-0.

[47] Neto E de M, da Silva EA, Nunes HR de C, Bazan R, de Souza LAPS, Luvizutto GJ. Effect of transcranial direct current stimulation in addition to visuomotor training on choice reaction time and cognition function in amateur soccer players (FAST trial): A randomized control trial. Neurosci Lett 2022;766:136346. 10.1016/j.neulet.2021.136346.

[48] Nguemeni C, Stiehl A, Hiew S, Zeller D. No Impact of Cerebellar Anodal Transcranial Direct Current Stimulation at Three Different Timings on Motor Learning in a Sequential Finger-Tapping Task. Front Hum Neurosci 2021;15:631517. 10.3389/fnhum.2021.631517.

[49] Orcioli-Silva D, Islam A, Baker MR, Gobbi LTB, Rochester L, Pantall A. Bi-Anodal Transcranial Direct Current Stimulation Combined With Treadmill Walking Decreases Motor Cortical Activity in Young and Older Adults. Front Aging Neurosci 2021;13:739998. 10.3389/fnagi.2021.739998.

[50] Parma JO, Profeta VL da S, Andrade AGP de, Lage GM, Apolinário-Souza T. TDCS of the Primary Motor Cortex: Learning the Absolute Dimension of a Complex Motor Task. J Mot Behav 2021;53:431–44. 10.1080/00222895.2020.1792823.

[51] Saimpont A, Mercier C, Malouin F, Guillot A, Collet C, Doyon J, et al. Anodal transcranial direct current stimulation enhances the effects of motor imagery training in a finger tapping task. Eur J Neurosci 2016;43:113–9. 10.1111/ejn.13122.

[52] Salvador MG, Ugrinowitsch H, Romano-Silva MA, Miranda DMD, Apolinário-Souza T, Lage GM. TRANSCRANIAL DIRECT CURRENT STIMULATION AND MANUAL ASYMMETRIES: THE EFFECT OF THE STIMULATION ON THE MANUAL DEXTERITY. J Phys Educ 2017;28. 10.4025/jphyseduc.v28i1.2837.

[53] Sevilla-Sanchez M, Hortobágyi T, Fogelson N, Iglesias-Soler E, Carballeira E, Fernandez-del-Olmo M. Small Enhancement of Bimanual Typing Performance after 20 Sessions of tDCS in Healthy Young Adults. Neuroscience 2021;466:26–35. 10.1016/j.neuroscience.2021.05.001.

[54] Talimkhani A, Abdollahi I, Mohseni-Bandpei MA, Ehsani F, Khalili S, Jaberzadeh S. Differential Effects of Unihemispheric Concurrent Dual-Site and Conventional tDCS on Motor Learning: A Randomized, Sham-Controlled Study. Basic Clin Neurosci 2019;10:59–72. 10.32598/bcn.9.10.350.

[55] von Rein E, Hoff M, Kaminski E, Sehm B, Steele CJ, Villringer A, et al. Improving motor performance without training: the effect of combining mirror visual feedback with transcranial direct current stimulation. Journal of Neurophysiology 2015;113:2383–9. 10.1152/jn.00832.2014.

